# PCK2 opposes mitochondrial respiration and maintains the redox balance in starved lung cancer cells

**DOI:** 10.1101/2020.11.23.393686

**Authors:** Gabriele Grasmann, Mélanie Planque, Corina T. Madreiter-Sokolowski, Andelko Hrzenjak, Wolfgang F. Graier, Sarah-Maria Fendt, Horst Olschewski, Katharina Leithner

## Abstract

Cancer cells frequently lack nutrients like glucose, due to insufficient vascular networks. A decrease of extracellular glucose is accompanied by enhanced mitochondrial respiration in cancer cells, which promotes the formation of potentially harmful reactive oxygen species (ROS). Here we show that a gluconeogenesis enzyme, mitochondrial phosphoenolpyruvate carboxykinase, PCK2, acts as a regulator of mitochondrial respiration and maintains the redox balance in nutrient-deprived lung cancer cells. PCK2 silencing increased the abundance and interconversion of tricarboxylic acid (TCA) cycle intermediates, augmented mitochondrial respiration and enhanced glutathione oxidation under glucose and serum starvation, in a PCK2 re-expression reversible manner. Moreover, augmenting the TCA cycle by PCK2 inhibition severely reduced colony formation. As a conclusion, PCK2 contributes to maintaining a reduced glutathione pool upon starvation besides mediating the biosynthesis of gluconeogenic/glycolytic intermediates. The study sheds light on adaptive responses in cancer cells to nutrient deprivation and identifies gluconeogenesis as starvation-induced pathway that limits respiration-induced oxidative stress.

## Introduction

Cancer cells undergo metabolic reprogramming for fast growth and proliferation. They utilize large amounts of glucose for the biosynthesis of cellular building blocks (Schulze & Harris, 2012; Vander Heiden *et al*, 2009). Moreover, certain amino acids as glutamine are consumed at high rates in order to support anabolic metabolism (DeBerardinis & Cheng, 2010; Schulze & Harris, 2012; Vander Heiden *et al*, 2009). Despite the induction of angiogenesis at an early stage of tumor growth, the nutrient supply is often not sufficient and steep nutrient gradients occur with increasing distance from the vessels (Vaupel, 2004). In a murine pancreatic cancer model, glucose levels were much lower in the tumor’s interstitial fluid than in the plasma, while glutamine levels remained unchanged (Sullivan *et al*, 2019). Thus, cancer cells need to adapt to a highly variable nutrient supply and starvation conditions (Cairns *et al*, 2011; DeBerardinis & Chandel, 2016).

As part of their metabolic rewiring, cancer cells are known to limit the complete catabolism of glucose, its entry into the TCA cycle and the rate of mitochondrial respiration by reducing the rate of acetyl-CoA formation at the step of pyruvate dehydrogenase (PDH) (Chandel, 2015). This key metabolic enzyme is repressed by pyruvate dehydrogenase kinase 1 (PDK1), which is induced by oncogenes like c-myc or β-catenin (Pavlova & Thompson, 2016), but also tightly regulated by allosteric inhibition by ATP and NADH (Chandel, 2015). In addition to PDH inhibition other regulatory mechanisms exist to balance TCA cycle activity and mitochondrial respiration. Acetyl-CoA is condensed with oxaloacetate (OAA) in the TCA cycle to form citrate, which eventually results in complete oxidation of acetyl-CoA to two CO_2_ molecules and subsequent regeneration of OAA. The concentration of OAA and acetyl-CoA are important regulators of the TCA cycle which fuels mitochondrial respiration by its production of reducing equivalents (Chandel, 2015; Krebs, 1970). In some conditions, including the high metabolic activity of cancer cells, TCA cycle intermediates get depleted due to their use in anabolic biosynthetic reactions (cataplerosis). Thus, anaplerosis, the replenishment of TCA cycle carbon e.g. from glutamate, is necessary to maintain the function of the TCA cycle (Owen *et al*, 2002). Cataplerosis, leading to reduced OAA availability, potentially limits mitochondrial respiration, especially in the context of glucose deprivation.

Glucose deprivation increases mitochondrial respiration in many cancer cell lines (Birsoy *et al*, 2014). However, mitochondrial respiration might enhance the formation of potentially damaging reactive oxygen species (ROS) as a certain amount of oxygen consumed by mitochondria is transformed to superoxide anion (O_2_^.-^), (Chandel, 2015). Superoxide is rapidly reduced to H_2_O_2_ by mitochondrial superoxide dismutase. H_2_O_2_ is a signaling molecule, but when produced in access it can give rise to the extremely reactive and cell damaging hydroxyl radical (OH·) upon reaction with metal cations (Fe^2+^ or Cu^+^). Thus, H_2_O_2_, is constantly converted to water by catalase, peroxiredoxins or by glutathione (GSH) –dependent enzymes including glutathione peroxidase 1 (GPX1), thereby oxidizing GSH to glutathione disulfide (GSSG) (Reczek & Chandel, 2015). GSSG is reduced again by glutathione reductase (GSR) using electrons from NADPH (Reczek & Chandel, 2015). The redox-modulated transcription factor nuclear factor erythroid 2-related factor 2 (NRF2, encoded by *NFEL2)* promotes antioxidant responses, among others by up-regulation of GSH synthesis via enhanced cystine import (Harris & DeNicola, 2020; Sasaki *et al*, 2002).

Phosphoenolpyruvate carboxykinase (PEPCK), a key enzyme of gluconeogenesis, catalyzes the GTP-dependent conversion of OAA to the glycolytic intermediate phosphoenolpyruvate (PEP). The enzyme exists in two different isoforms, the cytoplasmic isoform PCK1 (PEPCK-C), and the mitochondrial isoform PCK2 (PEPCK-M). Contrary to the previously assumed restriction to classic gluconeogenic tissues, PCK2 is also expressed in a variety of cancers, including lung cancer, breast cancer and prostate cancer (Chaika *et al*, 2012; Chen *et al*, 2007; Chu *et al*, 2017; Chun *et al*, 2010; Leithner *et al*, 2015; Leithner *et al*, 2018; Mendez-Lucas *et al*, 2014; Vincent *et al*, 2015; Zhao *et al*, 2017). However, PCK2 is also expressed in non-neoplastic tissues, including the lung (Smolle *et al*, 2020; Stark & Kibbey, 2014). The enzyme allows cancer cells to generate glycolytic intermediates from small non-carbohydrate molecules such as glutamine or lactate (reviewed in (Grasmann *et al*, 2019)). PCK2, the prime isoform expressed in lung cancer promotes the survival and proliferation of lung cancer cells under conditions of low glucose, as well as xenograft growth *in vivo* (Leithner *et al*, 2015; Leithner *et al*, 2018; Vincent *et al*, 2015). In case of low glucose availability, PCK2 mediates the biosynthesis of serine, glycine and purine nucleotides (Keshet *et al*, 2020; Vincent *et al*, 2015), or the glycerol backbone of phospholipids (Leithner *et al*, 2018) in cancer cells.

It remains unknown, whether PCK2 also regulates TCA cycle flux in cancer cells under nutrient starvation. Here we show that PCK2 diminishes the levels of TCA cycle intermediates in nutrient deprived lung cancer cells, thereby suppressing starvation-induced mitochondrial respiration. Moreover, we reveal that this cataplerotic activity protects lung cancer cells from growth inhibition by oxidative stress.

## Results

### PCK2 suppresses TCA cycle activity and limits TCA cycle intermediate abundance

The TCA cycle provides reducing equivalents to the respiratory chain. We assessed the abundance of TCA cycle intermediates and traced their interconversion in non-small cell lung cancer (NSCLC) cells by using uniformly ^13^C-labeled glutamine, the most important precursor for TCA cycle intermediates (DeBerardinis & Cheng, 2010; DeBerardinis & Chandel, 2016). To mimic conditions of full nutrient availability, the medium was supplemented with 10 mM glucose and 10% dialyzed fetal calf serum (dFCS). In contrast, a low concentration of glucose (0.2 mM) in serum-free medium was used for experiments under starvation conditions. Serum was omitted in starvation media, since it contains lipids and other macromolecular nutrients. Under starvation conditions, H23 and A549 lung cancer cells, showed a moderate decrease in the TCA cycle intermediates fumarate or malate, and a decline in the total amount of citrate, compared to non-starvation conditions (Fig 1C; Appendix Fig 1C). Glutamine provided carbons to TCA cycle intermediates under both conditions, leading to the full ^13^C labeling of malate and fumarate (denoted as M+4) (Fig 1A, B; Appendix Fig 1A, B). Accordingly, citrate M+4 was generated from the condensation of fully labeled OAA with unlabeled acetyl-CoA. Upon treatment with starvation media, citrate M+6 was formed from OAA (M+4) and fully labeled acetyl-CoA (M+2), which was very low under non-starvation conditions (Fig 1A, B; Appendix Fig 1A, B). This indicates a higher rate of conversion of TCA cycle metabolites via pyruvate to acetyl-CoA under treatment with starvation which has been already described to occur in absence of glucose (Yang *et al*, 2014a), mediated either via PEPCK or malic enzyme (ME). Accordingly, a significant proportion of pyruvate, the precursor of acetyl-CoA, was fully ^13^C labeled (M+3) in starvation, but not in non-starvation medium (Fig 1A, B; Appendix Fig 1A, B). A scheme of possible labeling patterns of TCA cycle intermediates, after the addition of ^13^C_5_-glutamine, is depicted in Fig 1E.

**Fig 1.**
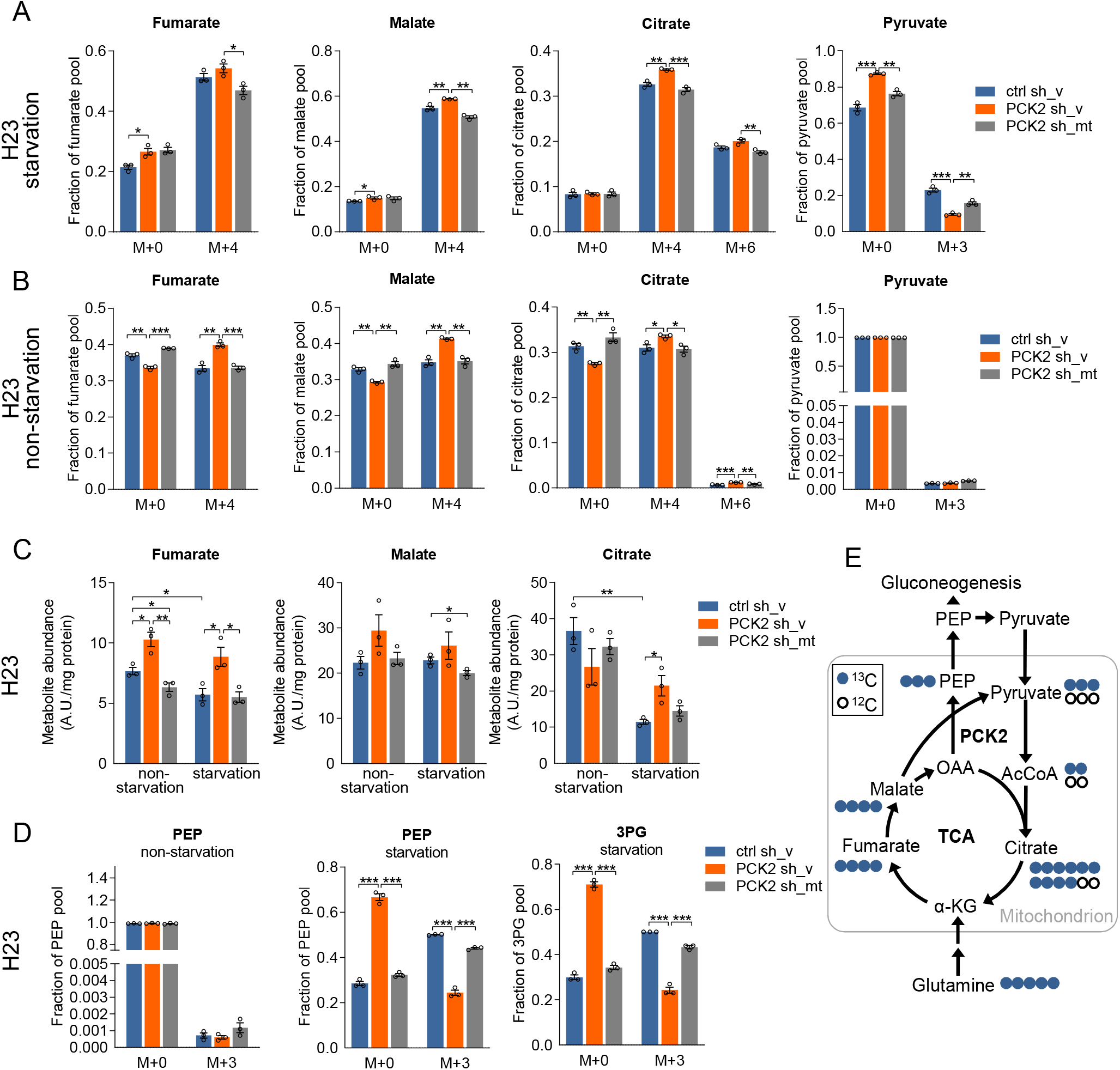
PCK2 silencing enhances stable isotopic labeling and abundance of TCA cycle intermediates and reduces labeling of downstream gluconeogenesis products. A, B, D H23 cells stably expressing non-silencing shRNA (ctrl sh) or PCK2 shRNA (PCK2sh) were transfected with the empty pCMV-AC vector (ctrl sh_v and PCK2 sh_v) or with a shRNA-resistant PCK2 allele (PCK2 sh_mt). Cells were treated for 24 hours with ^13^C_5_-glutamine and incorporation of ^13^C into TCA cycle intermediates was measured by GC-MS. Fractional enrichment of unlabeled (M+0) and fully labeled isotopologues in cells treated with starvation medium (A, D) (0.2 mM glucose, serum-free) or non-starvation medium (B, D) (10 mM glucose, 10% dialyzed FCS) containing ^13^C_5_-glutamine are shown. Results are shown as mean ± SEM from n=3 experiments. Group comparisons were made using two-sided, unpaired Student’s t-tests.* p< 0.05, ** p<0.01, *** p<0.001. PEP, phosphoenolpyruvate; 3PG, 3-phosphoglycerate. C Total abundances of TCA cycle intermediates in H23 cells treated with non starvation medium or starvation medium as described above. Results are shown as mean ± SEM from n=3 experiments. Group comparisons were made using twosided, unpaired Student’s t-tests* p< 0.05, ** p<0.01. A.U., arbitrary units. E Metabolic pathway for the conversion of ^13^C5-glutamine into TCA cycle and gluconeogenesis intermediates.

In order to address the role of PCK2 in tuning the TCA cycle under starvation conditions, PCK2 was silenced by stable expression of PCK2 shRNA (PCK2 sh). PCK2 knock-down was rescued by the expression of a point mutated, PCK2 shRNA resistant allele (PCK2 sh_mt; Appendix Fig 2B). In A549 cells, PCK2 was silenced by two different siRNA pools (Appendix Fig 2A). When total levels of TCA cycle intermediates were assessed, we found that PCK2 silencing clearly increased the levels of fumarate, malate and citrate under starvation conditions in both cell lines, in H23 cells the effect was blunted by the re-expression of shRNA resistant PCK2 (Fig 1C; Appendix Fig 1C). In H23 cells, M+4 labeling of citrate and malate was slightly increased by PCK2 silencing (Fig 1A). The effects were partly also observed at the level of fumarate. Likewise, silencing of PCK2 in A549 cells led to an increased abundance of M+4 isotopologues of malate and fumarate under starvation conditions (Appendix Fig 1A). In H23 cells, the fraction of pyruvate M+3 was decreased by PCK2 silencing (Fig 1A), while it was slightly enhanced in A549 cells (Appendix Fig 1A). These results indicate that PCK2 contributes to OAA decarboxylation in H23 cells, but not in A549 cells. The latter may utilize a different route of TCA cycle carbon to pyruvate conversion, e.g. via ME. Together, these data demonstrate that PCK2 removes OAA from the TCA cycle under starvation conditions, leading to a reduced abundance and interconversion of TCA cycle intermediates.

Importantly, the initial steps in glutamine carbon utilization, the conversion of glutamine to glutamate and the further transamination to α-ketoglutarate (α-KG), remained unaffected by PCK2 silencing when treated with starvation media (Appendix Fig 3A, B). Under non-starvation conditions, the fraction of fully labeled glutamate and α-KG (M+5), M+4 fumarate, malate and citrate and also the absolute abundance of α-KG and fumarate were increased by PCK2 silencing in H23 (Fig 1B, C; Appendix Fig 3A, C) but only partially in A549 cells (Appendix Fig 1B; Appendix Fig 3B, D). This may be related to a slightly, but not significantly enhanced expression of the initial enzyme in glutamine utilization, glutaminase *(GLS1)* and a slight decrease in the expression of the cataplerotic enzyme ATP citrate lyase *(ACLY)* in H23 cells upon PCK2 silencing under non-starvation conditions (Appendix Fig 3E).

PCK2 contributed to the gluconeogenesis pathway under starvation conditions, as shown by the high rate of conversion of ^13^C_5_-glutamine via OAA to PEP. Between 40 and 50% of PEP showed a full labeling by ^13^C and a similar enrichment was found at the level of the downstream gluconeogenesis intermediate 3-phosphoglycericate (3PG) (Fig 1D; Appendix Fig 1D). Only a small fraction of PEP (0.1 to 1%) was labeled under treatment with non-starvation media, showing that PCK2 activity in the direction of gluconeogenesis was low under these conditions. This was accompanied by a reduced expression of PCK2 (Appendix Fig 2A), similar to our previous findings (Leithner *et al*, 2015; Leithner *et al*, 2018). Importantly, PCK2 silencing decreased the fraction of ^13^C-labeled PEP and 3PG (Fig 1D; Appendix Fig 1D). In A549 cells, starvation treatment increased the abundance of PEP whereas it was reduced by PCK2 silencing (Appendix Fig 1D).

### PCK2 decreases mitochondrial respiration under starvation conditions

The TCA cycle produces reducing equivalents, which are oxidized in the electron transport chain (ETC) to generate ATP (Martínez-Reyes & Chandel, 2020). When we measured oxygen consumption rates (OCR), we found an increase in mitochondrial respiration under starvation compared to non-starvation conditions (Fig 2A-G). Starvation-induced basal respiration was further clearly enhanced upon silencing of PCK2 (Fig 2B-G). In both cell lines, PCK2 silencing increased both, basal and maximal OCR, but not ATP-linked respiration under treatment with starvation media, suggesting that the additional oxygen consumed under PCK2-silenced conditions is not utilized for ATP biosynthesis (Fig 2D-G). No significant effect of PCK2 silencing was found under nonstarvation conditions (Fig 2D-G). During OCR measurements either pyruvate or lactate were added as a respiratory fuel. Of note, the cells respired also in the absence of pyruvate or lactate (starvation-lac, Fig 2B), with a similar enhancement by PCK2 silencing compared to cells under starvation media supplemented with lactate. Treatment with etomoxir (Eto), an inhibitor of fatty acid oxidation, decreased the level of OCR under starvation conditions, indicating that in the absence of glucose and serum, cells partially utilize (endogenous) fatty acids to fuel respiration (Fig 2C). Also if fatty oxidation was blocked, PCK2 silenced cells showed elevated oxygen consumption rates (Fig 2C). These data indicate that mitochondrial respiration is decreased by PCK2.

**Fig 2.**
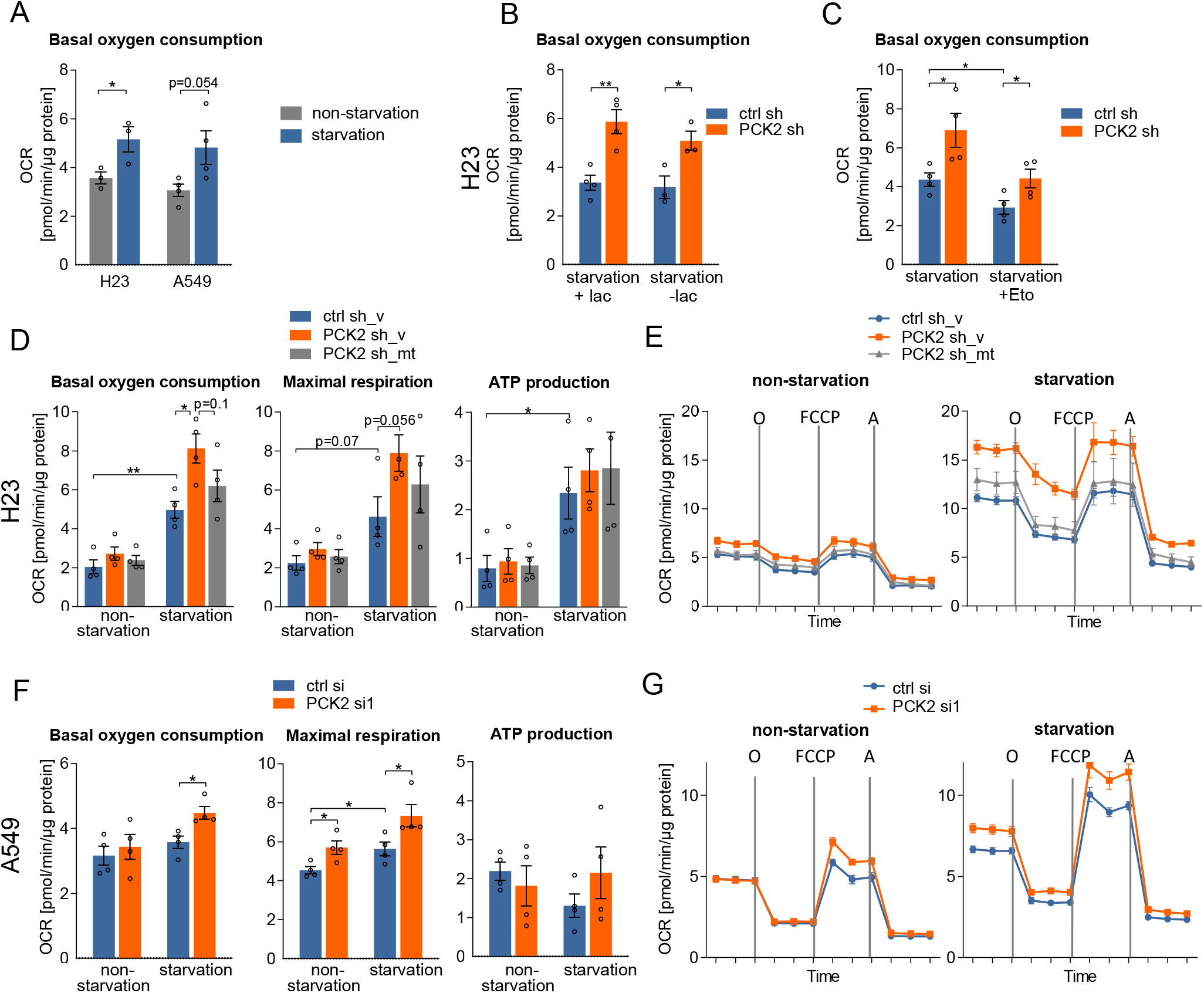
PCK2 silencing enhances cellular respiration. A Basal oxygen consumption rate (OCR) in H23 or A549 cells treated for 24 hours with non-starvation medium (10 mM glucose and 10% dialyzed serum) or starvation medium (0.2 mM glucose, serum-free). B, C H23 cells stably expressing non-silencing shRNA (ctrl sh) or PCK2 shRNA (PCK2 sh) were cultured for 24 hours in starvation medium and the OCR was measured in the presence or absence of lactate (B) or etomoxir (Eto), a fatty acid oxidation inhibitor (C). D, E Basal and maximal OCR and ATP-production-linked OCR in H23 cells stably expressing non-silencing shRNA (ctrl sh) or PCK2 shRNA (PCK2 sh) and transfected with the empty pCMV-AC vector (ctrl sh_v and PCK2 sh_v) or with a shRNA-resistant PCK2 allele (PCK2 sh_mt), followed by treatment with starvation or non-starvation medium. (E) Representative OCR recordings over time at basal levels and after the addition of oligomycin (O), FCCP or antimycin (A). F, G A549 cells were transfected either with non-silencing ctrl siRNA (ctrl si) or with PCK2 targeting siRNA (PCK2 si1) and treated as described above. A-D, F Results are shown as mean ± SEM from four independent experiments, each consisting of four to six technical replicates. Comparison between groups was performed using two-sided, unpaired Student’s t-tests.* p< 0.05, ** p<0.01. E, G Representative recordings showing mean values ± SEM of five or six technical replicates.

Mitochondrial morphology can vary under different nutritional conditions. Mitochondria tend to be elongated in case of inappropriate nutrient supply as this protects them from autophagosomal degradation and induces increased oxidative phosphorylation (OXPHOS) (Mishra & Chan, 2016; Rambold *et al*, 2011). When mitochondrial morphology was visualized by Mitotracker green, the number of individual mitochondria did not differ between starvation and non-starvation conditions (Appendix Fig 4A). However, PCK2 silencing decreased the number of individual mitochondria under starvation conditions, linking PCK2 activity to a decrease in mitochondrial elongation (Appendix Fig 4A). Mitochondrial mass was not significantly affected, neither by treatment with starvation media nor by PCK2 silencing (Appendix Fig 4B). In order to clarify, whether an upregulated expression of complex members of the respiratory chain causes enhanced mitochondrial respiration under PCK2 silencing, we assessed the protein abundance of key electron transport chain subunits. The expression of NDUFB8 (complex I), SDHB (complex II), UQCRC2 (complex III), COX II (complex IV) and ATP5A (F1 subunit of complex V) and the mitochondrial protein TOM20 remained unchanged (Appendix Fig 4C, D). Thus, enhancement of respiration by PCK2 silencing occurs rather due to modulation of the TCA cycle activity than due to altered expression of OXPHOS members.

### PCK2 improves the redox balance in starved lung cancer cells

The respiratory chain is the major source of potentially harmful ROS which need to be continuously scavenged by different antioxidant enzymes at the expense of NADPH and GSH (Chandel, 2015; Harris & DeNicola, 2020; Reczek & Chandel, 2015). We found that *NFEL2* and different antioxidant enzymes utilizing or providing GSH, including *GSR* and *GPX4,* the cysteine transporter subunit *SLC7A11* and the uncoupling protein *UCP2* were up-regulated in NSCLC cells after 24 hours of treatment with starvation media (Appendix Fig 5). PCK2 silencing led to a slight but significant suppression of starvation-induced SLC7A11 expression (Appendix Fig 5).

Interrogating, if the increased activity of the mitochondrial respiration, observed under starvation conditions and triggered by PCK2 silencing, leads to an enhanced formation of ROS, we measured mitochondrial superoxide and cellular ROS. Treatment with starvation media resulted in a slight increase in mitochondrial superoxide, which was not significantly altered by PCK2 silencing (Fig 3A). Likewise, DCFDA oxidation, a marker of increased ROS levels, showed only a small, insignificant increase under starvation conditions and PCK2 silencing (Fig 3B). However, PCK2 silencing, under starvation conditions significantly decreased the ratio of reduced to oxidized GSH (GSH/GSSG), as shown by two different methods (Fig 3C, Appendix Fig 6A). Moreover, the NADPH/NADP^+^ ratio was decreased upon PCK2 silencing under treatment with starvation media (Fig 3D).

**Fig 3.**
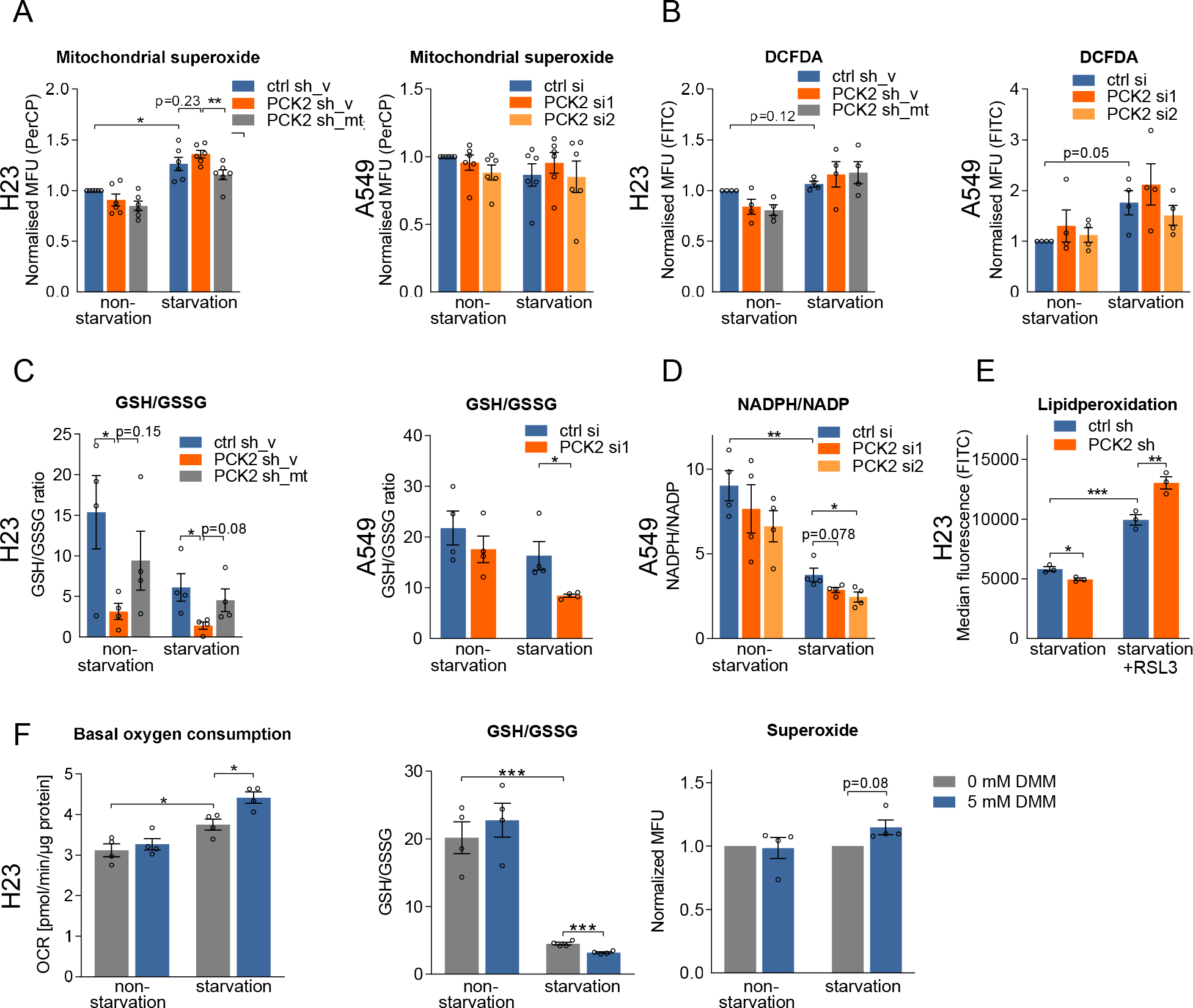
PCK2 silencing substantially decreases the ratio of reduced to oxidized glutathione, dimethyl-L-malate addition mimics PCK2 silencing. A Mitochondrial superoxide measured by MitoSOX in H23 cells stably expressing ctrl sh or PCK2 sh and empty pCMV-AC vector (H23 ctrl sh_v and PCK2 sh_v) or a PCK2 shRNA resistant PCK2 allele (PCK2 sh_mt) and in A549 cells, transfected with non-silencing (ctrl si) or PCK2 silencing siRNA (PCK2 si1/2) and treated with non-starvation or starvation medium for 24 hours. Results are shown as mean ± SEM from n=6 experiments. B Oxidation of CM-H_2_DCFDA, an indicator of cellular H_2_O_2_ and other reactive oxygen species in cells treated as described in (A). Results are shown as mean ± SEM from n=4 experiments. C Reduced (GSH) to oxidized (GSSG) glutathione ratio measured by a commercially available kit in cells treated as described in (A). Results are shown as mean ± SEM from n=4 experiments. D Ratio of NADPH to NADP measured by liquid chromatography coupled to tandem mass spectrometry (LC-MS/MS) in cells treated as described in (A). Results are shown as mean ± SEM from n=4 experiments. E Lipid peroxidation in H23 cells treated with starvation medium with or without glutathione peroxidase 4 inhibitor RSL3. Results are shown as mean ± SEM from n=3 experiments. F H23 cells were treated with non-starvation or starvation media containing 5 mM dimethyl-L-malate (DMM), a cell-permeable malate analogue. Basal oxygen consumption, GSH/GSSG ratio and mitochondrial superoxide production were measured. Results are shown as mean ± SEM from n=4 experiments. A-F Comparison between groups was performed using two-sided, unpaired Student’s t-tests or one group Student’s t-test as applicable.* p< 0.05, ** p<0.01, *** p<0.001.

Lipid peroxidation is a self-perpetuating detrimental oxidative process that may lead to a specific, iron dependent, form of cell death, ferroptosis (Harris & DeNicola, 2020). In cancer cells, lipid peroxidation is efficiently controlled by the lipid specific enzyme glutathione peroxidase 4 (GPX4) which reduces lipid peroxidation by utilizing reduced GSH (Yang *et al*, 2014b). If the GPX4 antioxidant system was blocked by the GPX4 inhibitor RSL3, PCK2 silencing caused significantly enhanced lipid peroxidation levels, indicating a higher burden of ROS and/or an insufficiency of alternative antioxidant defense mechanisms (Fig 3E). An enhanced expression of antioxidant enzymes under starvation conditions and a decreased GSH/GSSG ratio suggest that ROS formed by the electron transport chain under PCK2 silencing may be scavenged by antioxidant defense mechanisms at the expense of a diminished glutathione redox capacity. In H23 cells, but not A549 cells, GSH depletion induced by PCK2 silencing was also observed under non-starvation conditions (Fig 3C). Oxygen consumption was only insignificantly increased under these conditions (Fig 2D, E), despite the enhanced abundance of TCA cycle intermediates in this cell line (Fig 1C). Potentially, ROS formation from enhanced TCA cycle intermediates, independent of a forward oxidation in the respiratory chain, might play a role, e.g. succinate-fueled reverse-electron flow (Chouchani *et al*, 2014).

In order to clarify whether increased TCA activity and subsequently enhanced respiration were responsible for the GSH/GSSG imbalance, we utilized dimethyl-L-malate (DMM), a membrane permeable analogue of the TCA cycle intermediate malate. Fueling the TCA cycle with 5 mM DMM mimicked the effects of PCK2 silencing on respiration and depleted GSH in H23 cells, while not increasing superoxide (Fig 3F). Partly, these effects were also observed in A549 cells (Appendix Fig 7A-C). These data show that PCK2 activity diminishes oxidative stress and contributes to maintaining the GSH/GSSG redox balance by reducing respiration via suppression of the TCA cycle.

GSH levels are regenerated by the reduction of GSSG, however, GSH levels are also maintained by *de novo* synthesis from glycine, cysteine and glutamate (Bansal & Simon, 2018). Interestingly, PCK2 silencing was associated with a slightly reduced rate of GSH *de novo* synthesis, as shown by the lower fractional abundance of GSH M+5, in starved H23 but not A549 cells (Appendix Fig 6B). The M+5 labeled GSH found under starvation conditions likely reflects labeled glutamate (carrying 5 carbons), which is directly formed from ^13^C-glutamine. We could not detect a transfer of ^13^C from glutamine to serine/glycine or cysteine in H23 cells and a low level of transfer in A549 cells, under our experimental conditions (data not shown). Accordingly, we did not detect higher isotopologues in the GSH pool under these experimental conditions. These data indicate that PCK2 slightly promotes GSH *de novo* biosynthesis in H23 cells independent of glycine biosynthesis.

### PCK2 promotes colony formation under starvation conditions and reduces the sensitivity towards H_2_O_2_

Next, we investigated colony formation by lung cancer cells under starvation or non-starvation conditions. After an initial starvation or non-starvation treatment, we allowed cells to recover and form colonies in full growth medium. As reported previously (Leithner *et al*, 2018), PCK2 silencing significantly diminished colony formation under starvation (Fig 4A, B). In order to investigate whether the reduced colony formation by PCK2 silencing is caused by the redox imbalance, we added different antioxidants simultaneously with starvation or non-starvation treatment. Trolox, a derivative of vitamin E, exogenous GSH, as well as the antioxidant and GSH precursor N-acetyl cysteine (NAC) all rescued the effect of reduced colony forming ability in PCK2 silenced cells (Fig 4A, B). Importantly, enhancing TCA cycle intermediates by using the malate analogue DMM mimicked the impact of PCK2 silencing on colony formation in H23 but not in A549 cells (Fig 4C; Appendix Fig 7D). The generally less pronounced effects of DMM supplementation in A549 cells might be attributed to a higher efflux/decarboxylation of TCA intermediates through ME in that cell line, which generates mitochondrial NADPH. Additionally, we investigated whether PCK2 expression protects starved cancer cells from damage induced by exogenous H_2_O_2_. In both cell lines, PCK2 silencing significantly enhanced the toxic effects of H_2_O_2_ under treatment with starvation (Fig 4D). No effect of PCK2 silencing on cell numbers or proliferation was found in densely seeded cells under starvation conditions (Fig 4D, E). This finding is in striking contrast to the reduction of colony formation by PCK2 in starvation conditions. In order to clarify, whether colony forming cancer cells are more sensitive towards oxidation than densely seeded cells, we used butionine sulfoximine (BSO), an inhibitor of GSH biosynthesis and inducer of oxidative stress. In fact, treatment with different concentrations of BSO highly reduced the cellular colony forming ability upon non-starvation and starvation conditions, whereas it affected cell numbers in densely seeded cells to a much lower extent (Appendix Fig 8).

**Fig 4.**
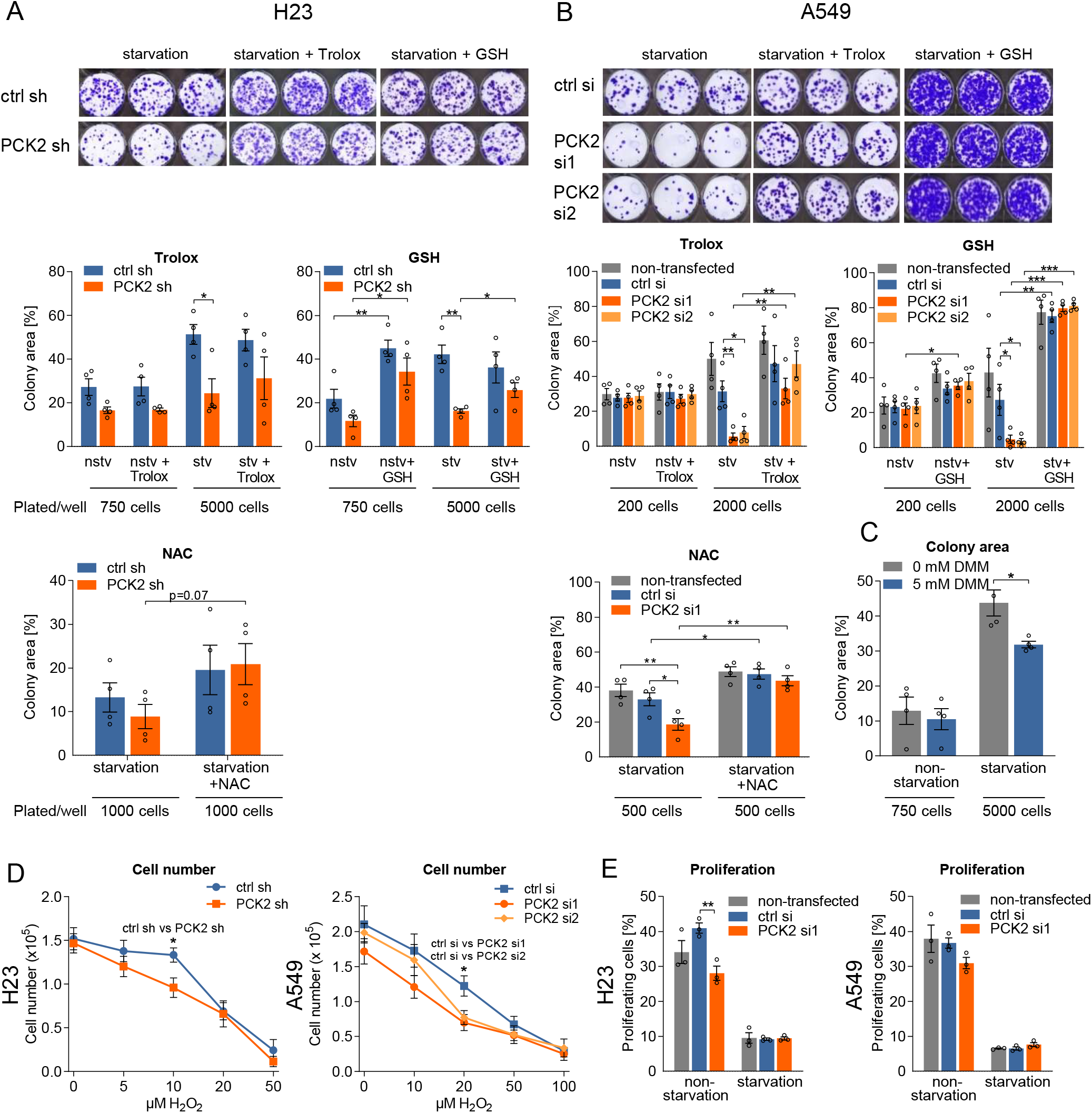
PCK2 silencing reduces colony forming capability of H23 and A549 cells, rescued by antioxidants, phenocopy by dimethylmalate. A H23 cells expressing control shRNA or a PCK2 shRNA were plated at suitable densities for colony formation and treated with non-starvation (nstv) or starvation (stv) media for 72 hours, followed by a recovery in normal growth media. The antioxidants Trolox (100 μM), glutathione (GSH, 2 mM), or N-acetyl cysteine (NAC, 10 mM) were added simultaneously with nstv or stv treatment. Assays were performed in technical triplicates. Data are mean ± SEM from n=4 independent experiments. B A549 cells were non-transfected or transfected with non-silencing (ctrl si) or PCK2 silencing siRNA (PCK2 si1/2) and treated as above with or without Trolox (100 μM), GSH (2 mM), or NAC (5 mM). Data are mean ± SEM from n=4 independent experiments. C H23 cells were plated and treated with nstv/stv media as described above. During the treatment period, 5 mM dimethyl-L-malate (DMM) was added. Data are mean ± SEM from n=4 independent experiments. D Cell counts of densely plated control or PCK2 silenced cells, treated with starvation media containing different concentrations of H_2_O_2_. The assay was performed in technical duplicates. Data are mean ± SEM from n=4 independent experiments. E EdU incorporation into densely plated control or PCK2 silenced cells, treated with non-starvation or starvation media. Data are mean ± SEM from n=3 independent experiments. A-E Group comparisons were made using two-sided, unpaired Student’s t-tests. * p< 0.05, ** p<0.01, *** p<0.001.

### Addition of the protein glutathionylation agent diamide mimics PCK2 silencing

GSSG-induced modifications, independent of direct cellular damage by ROS, might contribute to the suppressive effect of PCK2 silencing on colony formation. High GSSG and low GSH levels have been found to promote S-glutathionylation of proteins (Mieyal *et al*, 2008). To address the question whether protein S-glutathionylation may be responsible for PCK2 knockdown induced reduction of colony formation, we added diamide, an S-glutathionylating agent, to starvation or non-starvation media. Diamide concentration-dependently caused a decrease in the colony forming ability under treatment with starvation, but not non-starvation media, indicating that starvation conditions render colony forming cancer cells vulnerable towards glutathionylation of proteins (Fig 5A). Additionally, enzymes responsible for protein de-glutathionylation, such as, sulfiredoxin *(SRXN1)* and glutaredoxin-1 *(GLRX)* but not glutathione S-transferase P *(GSTP),* which selectively glutathionylates proteins, were up-regulated upon starvation conditions in both cell lines (Fig 5B).

**Fig 5.**
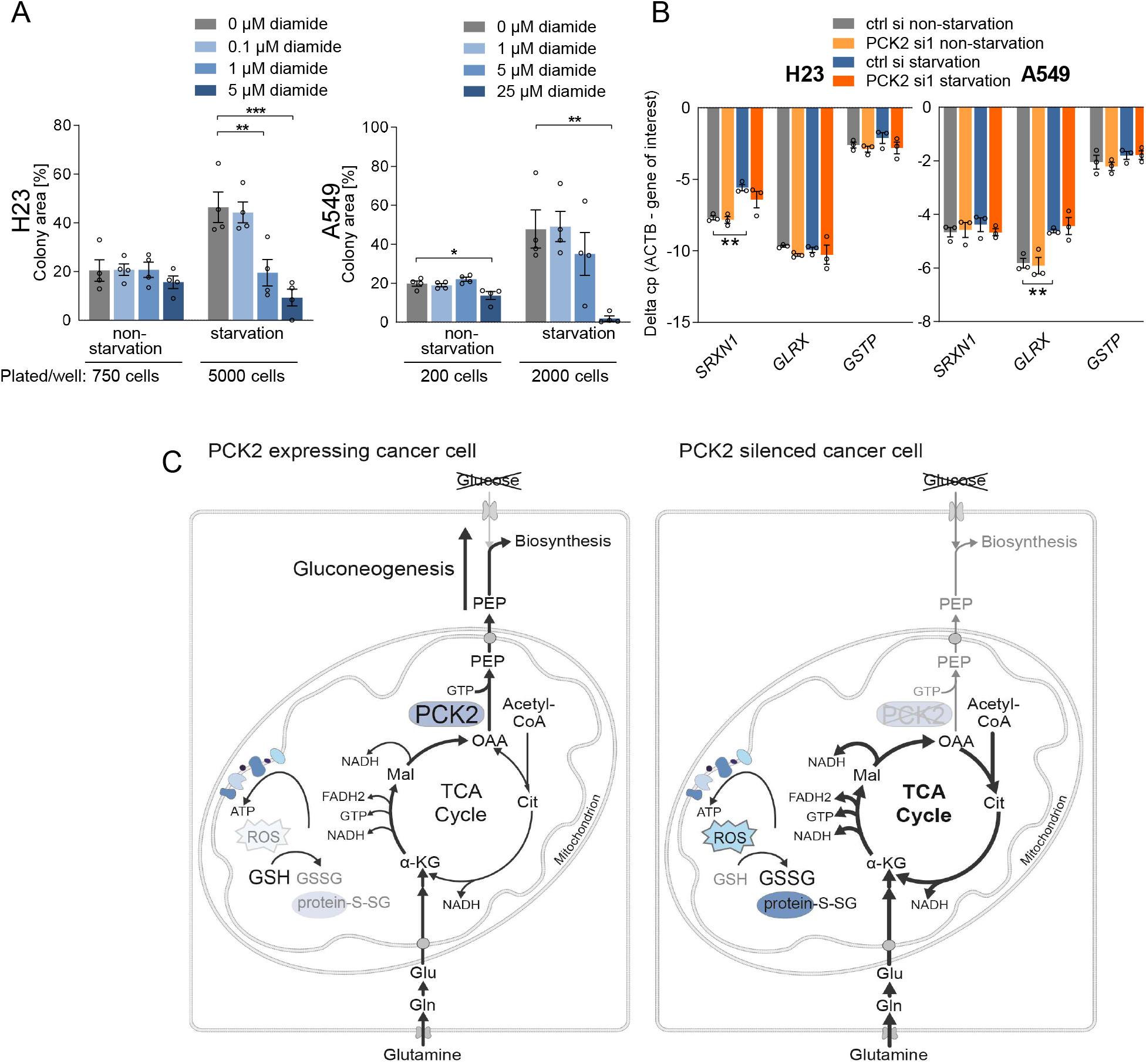
Diamide, a glutathionylating agent, phenocopies the effects of PCK2 silencing. A H23 and A549 cells were plated for colony formation assay and treated with non starvation or starvation media in the presence or absence of diamide. Group comparisons were made using one-way ANOVA and Dunnett post hoc analysis. Data are shown from n=4 independent experiments, each containing three technical replicates. * p< 0.05, ** p<0.01, *** p<0.001. B H23 or A549 cells were transfected either with non-silencing ctrl siRNA or with PCK2 silencing RNA (PCK2 si1) and treated for 24 hours with non-starvation or starvation media. Expression of genes (normalized to β-actin, ACTB) was analyzed by quantitative PCR. Results are shown as mean ± SEM from n=3 independent experiments. Group comparisons were made using two-sided Student’s t-tests. ** p<0.01. C Model for the cataplerotic activity of PCK2 in cancer cells, reducing respiration and maintaining a reduced glutathione redox potential, particularly under starvation conditions.

In summary, these results demonstrate that the cataplerotic action of PCK2 directly reduces TCA cycle intermediate abundance and interconversion, diminishes mitochondrial respiration and depletes cellular GSH. The rescue by antioxidants and the phenocopy by a protein glutathionylating agent suggest that PCK2-mediated maintenance of the GSH/GSSG balance is important during colony formation under nutrient starvation. A model for the protective role of PCK2 in mediating cataplerosis, limiting respiration and maintaining the glutathione redox balance in cancer cells is proposed in Fig 5C.

## Discussion

In many tumor types, the utilization of certain steps of gluconeogenesis is beneficial in starved cancer cells, since this pathway yields building blocks for biomass production (Grasmann *et al*, 2019). The TCA cycle is a metabolic hub, feeding into biosynthetic pathways for the generation of amino acids, nucleic acids or fatty acids, but the cycle also produces reducing equivalents, critical for electron transport chain and ATP production (Martínez-Reyes & Chandel, 2020; DeBerardinis & Cheng, 2010; Owen *et al*, 2002;). Here, we identify the importance of PCK2 as a regulator of the TCA cycle in lung cancer cells, subsequently limiting mitochondrial respiration and enhancing the antioxidant defense.

We show that PCK2 silencing causes increased abundance of TCA cycle intermediates, especially in low glucose, serum-free media. Accordingly, we found augmented mitochondrial respiration under starvation conditions, which got even enhanced by PCK2 silencing. Electron leakage from the ETC causes the formation of ROS, which are scavenged by antioxidant enzymes at the expense of GSH (Reczek & Chandel, 2015). Treatment with starvation media induced the antioxidant genes *NFEL2, SLC7A11, GSR* and *GPX4.* The enhancement of *NFEL2* and *SLC7A11* by starvation has been previously described (Koppula *et al*, 2017). Many antioxidant systems, including GPX4, utilize GSH as a co-factor (Reczek & Chandel, 2015). In fact, the GSH/GSSG ratio and consequently the NADPH/NADP^+^ ratio were decreased by PCK2 silencing under treatment with starvation media. ROS levels remained unchanged by PCK2 silencing under these experimental conditions, showing that ROS were continuously scavenged by the antioxidant systems, at the expense of reduced glutathione. However, when lipid peroxidation was initiated by addition of a GPX4 inhibitor, PCK2 silencing led to an increased level of oxidized phospholipids. In summary, these data demonstrate that PCK2 plays an important role in maintaining the cellular antioxidant defense in lung cancer upon treatment with starvation media. In addition to the prevention of the accumulation of free ROS, a decreased GSH/GSSG ratio provokes modifications of signaling proteins by S-glutathionylation e.g. protein kinase C or NF-κB, thereby altering their function (Mieyal *et al*, 2008). Of note, we found a highly increased expression of sulfiredoxin or GLRX, crucial, GSH-dependent deglutathionylating enzymes, under starvation conditions, possibly related to an enhanced rate of protein-glutathionylation.

The role of PCK2 in tumor cells, pro- or antitumorigenic, has been previously linked to its effects on the TCA cycle, although the role of PCK2 under glucose and serum starvation has not been addressed in these studies. Under high glucose conditions, PCK2 silencing has been found to increase TCA cycle intermediate abundance in tumor initiating cells of melanoma and prostate cancer, as well as in hypoxic breast cancer cells (Luo *et al*, 2017; Tang *et al;* Zhao *et al*, 2017). However, the effect of PCK2 on the incorporation of ^13^C_5_-glutamine into the TCA cycle was not analyzed. While TCA cycle suppression by PCK2 and the decrease in citrate were associated with enhanced proliferation in prostate cancer cells due to diminished protein acetylation and superoxide levels (Zhao *et al*, 2017), the same effect of PCK2 on the TCA cycle was linked to growth inhibition in melanoma cells (Luo *et al*, 2017). Interestingly, in breast cancer cells, inhibition of the TCA cycle and suppression of ROS formation by PCK2 overexpression appeared to cause a growth arrest (Tang *et al*). In glucose but not serum starved A549 cells, PCK2 silencing did not significantly alter glutamine derived anaplerosis of the TCA cycle (Vincent *et al*, 2015), however, in contrast to our study, medium containing 0 mM glucose and 10% dFCS was utilized and abundances of isotopologues were normalized to high glucose conditions. PCK1 overexpression in liver cancer cells was shown to reduce TCA metabolite levels due to cataplerosis by PEPCK in high and low glucose containing media, thereby enhancing ROS formation and reducing cell viability due to an energy crisis (Liu *et al*, 2018). In contrast, PCK1 silencing reduced the abundance of ^13^C_5_ glutamine or ^13^C_3_ lactate derived TCA cycle intermediates in colon cancer cells (Montal *et al*, 2015; Montal *et al*, 2019). Given the variable effects of PCK1/2 on TCA cycle in different studies and systems, the question arises, whether different rates of the TCA cycle in different tissues of origin may determine the overall effect of PCK1/2 on TCA cycle intermediate abundance and TCA cycle flux. In non-cancerous liver and intestine PCK1 knockout mice displayed an increase in TCA cycle intermediates connected to decreased cataplerosis (Burgess *et al*, 2004; Potts *et al*, 2018; Satapati *et al*, 2015; She *et al*, 2000). In contrast to our findings obtained in lung cancer cells, the cataplerotic effect of PCK1 rather enhanced than repressed TCA cycle flux and respiration in liver from mice fed a high fat diet (Satapati *et al*, 2015). This was attributed to a reduced feedback inhibition by NADH (Satapati *et al*, 2015). Thus, the effect of PCK1 and PCK2 might be context-dependent and related to the energy/NADH status of the mitochondria.

The multifaceted role of the TCA cycle as a metabolic hub that controls anabolic pathways and ROS production gains further complexity by the fact that it may promote the production of NADPH either directly by NADP^+^ utilizing isoforms of isocitrate dehydrogenase (e.g. IDH2), or indirectly by feeding intermediates into the ME pathway (DeBerardinis & Cheng, 2010; DeBerardinis & Chandel, 2016). In our study, we found that promoting the TCA cycle in lung cancer cells, either by PCK2 silencing or by exogenous addition of DMM, enhanced respiration and decreased the GSH/GSSG redox potential. The data are in line with a previous report on mitochondrial respiration enhanced by silencing of ME and a dose dependent enhancement of ROS formation by DMM in lung cancer cells (Ren *et al*, 2014). The up-regulation of antioxidant defense mechanisms, e.g. the induction of the NRF2 target gene *SLC7A11* by starvation conditions found in our study and in previous reports (Koppula *et al*, 2017), indicates that low nutrient supply is associated with a higher burden of ROS in cancer cells. Glucose deprivation as well as complete nutrient starvation mediated by Hank’s balanced salt solution (HBSS) treatment were shown to trigger ROS formation (De Saedeleer *et al*, 2014; Jelluma *et al*, 2006; Koppula *et al*, 2017; Owada *et al*, 2013). When lipid oxidation was measured in tumor-bearing mice fed with a ketogenic, low-carbohydrate diet, an increase in lipid peroxidation was found upon radiation therapy and supporting the efficacy of the treatment (Allen *et al*, 2013). The reason for enhanced burden of ROS under glucose deprivation conditions is not completely understood, however increased respiration or reduced regeneration/biosynthesis of antioxidant molecules, such as NADPH or GSH, might play a role.

Our study suggests an important role in TCA cycle cataplerosis by PCK2 to balance respiration and ROS formation under starvation conditions. Interestingly, PCK2 silencing did not enhance ROS formation and proliferation in cells growing in high confluency, but dramatically reduced colony formation under starvation conditions. We found that loosely seeded, colony forming cells are more susceptible towards oxidative stress compared to densely seeded cells. It has been shown previously that certain cell types have enhanced ROS levels upon loose seeding densities (Pani *et al*, 2002). In line with an enhanced dependency of colony forming cells on antioxidant systems, GSH and its precursor NAC did not only rescue decreased colony forming ability by PCK2 silencing but increased colony formation upon treatment with starvation media also in control cells. Importantly, both, the TCA cycle intermediate DMM and the protein glutathionylating agent diamide phenocopied the effects of PCK2 silencing. Together, these data demonstrate that the redox balancing effect of PCK2 is beneficial for survival and colony formation of starved lung cancer cells. The phenotypic changes in cancer cells underlying the inhibition of colony formation induced by PCK2 silencing and GSH-loss remain unclear and should be investigated in future studies. Besides ROS-induced cell death, a diminished proliferation due to alterations in cell signaling might play a role. It has been previously shown that PCK2 mediates the generation of glycerol phosphate for phospholipid backbone biosynthesis in glucose starved lung cancer cells and that exogenous phosphatidylethanolamine (PE) phospholipids partially rescue colony formation inhibited by PCK2 knockdown (Leithner *et al*, 2018). Of note, PE, especially when containing polyunsaturated fatty acids, is highly susceptible to peroxidation (Kagan *et al*, 2017). Thus, PE backbone synthesis and turnover, facilitated by PCK2, might act together with the TCA suppressing effects of PCK2 described here to reduce ROS induced alterations under starvation conditions.

We show that PCK2 diminishes the TCA cycle and reduces starvation-induced mitochondrial respiration in starved lung cancer cells, thus maintaining the glutathione redox balance. This balance, however, is crucial for colony formation capabilities especially under starvation conditions and protects cells from the attack by exogenous ROS. Thus, PCK2 inhibition represents a potential new therapeutic approach to prevent metabolic adaptation and distort the redox balance in lung cancer cells.

## Methods

### Cell lines

The human NSCLC cell line A549 was obtained from Cell Lines Service (Eppelheim, Germany). A549 cells were cultured in DMEM/F-12 (Gibco, Waltham, MA, USA) supplemented with 10% fetal calf serum (FCS, Biowest, Nuaillé, France) and antibiotics (Gibco, Waltham, MA, USA). The human NSCLC cell line NCI-H23 (H23) was purchased from American Type Culture Collection (ATCC, Manassas, VA, USA). H23 cells were cultured in RPMI 1640 (Gibco) supplemented with 10% FCS (Biowest, Nuaillé, France) and antibiotics (Gibco). If not stated differently, cells were plated for all experiments with a density of 22,000 cells per cm^2^ in normal growth media. After 20-24 hours cells were washed twice with phosphate buffered saline (PBS) and treated either with non-starvation or with starvation media. For starvation/non-starvation treatment, glucose and glutamine free DMEM (Gibco) or glucose and glutamine free RPMI SILAC medium (Gibco) were supplemented with 200 mM glutamine, 0.2/10 mM (starvation/non-starvation) glucose, 0%/10% dialyzed FCS and antibiotics. RPMI SILAC was additionally supplemented with 1.15 mM arginine and 0.27 mM lysine. Cell line authentication was done for both cell lines by Short Tandem Repeat (STR) analysis using the PowerPlex 16HS System (Promega, Madison, WI).

### Stable expression of PCK2 shRNA

H23 cells were stably transfected with PCK2 shRNA or non-silencing control shRNA (Qiagen, Hilden, Germany). Puromycin-selection and generation of monoclonal subcultures was performed as described (Leithner *et al*, 2018). Maintenance media was supplemented with 0.5 μg/μL puromycin (Sigma Aldrich, Waltham, MA, USA) which was omitted during experiments. To confirm the specificity of PCK2 silencing effects, we transfected PCK2 silenced cells with a shRNA resistant PCK2 cDNA construct. Three codons of the shRNA binding site were point mutated without altering the amino acid sequence (Eurofins Genomics, Ebersberg, Germany) and cloned into pCMV6-AC expression vector (Origene, Rockville, MD); the empty expression vector was used as a control. Cells were transfected with 0.5-1 μg DNA/200,000 cells 48 hours before treatment start with jetPRIME (Polyplus, Illkirch, France) transfection reagent.

### PCK2 silencing with siRNA

A549 and H23 cells were transfected either with non-silencing (ctrl) siRNA (Non-targeting pool, Dharmacon, Horizon, Colorado, USA) or with PCK2 siRNA1 (Smart pool PCK2, Dharmacon). PCK2 siRNA2 was a custom designed pool with the sequences GGAUGAGGUUUGACAGUGA and UGGCUACAAUCCAGAGUAA (Dharmacon).

### Stable isotopic tracing

For analysis of TCA cycle metabolite abundance and flux, ^13^C_5_-glutamine was utilized as a tracer. Cells were treated for 24 hours with glutamine free non-starvation/starvation media supplemented with 2 mM ^13^C_5_-glutamine. Then cells were washed with saline and metabolism was quenched by immediately freezing the cells on liquid nitrogen.

### Sample extraction and GC-MS and LC-MS measurements

Gas chromatography-mass spectrometry (GC-MS) and liquid chromatography-mass spectrometry (LC-MS) was performed essentially as described (Kampen *et al*, 2019; Lorendeau *et al*, 2017). For details see Appendix Methods.

### Seahorse measurements

Cells were plated on Cell-Tak™ coated XF96 polystyrene cell culture microplates (Seahorse Bioscience®, Agilent; California, US) at a density of 40,000 cells per well 24 hours before treatment. Thereafter, cells were cultured for 24 hours in starvation or nonstarvation media. As respiratory substrate we added either 1 mM pyruvate (Sigma) (experiment with H23 ctrl sh_v/PCK2sh_v and PCK2sh_mt cells) or 10 mM lactate (Sigma) (experiments with H23 ctrl sh/PCK2 sh and A549 cells), if not indicated differently. OCR was measured with a XF96 Extracellular Flux analyzer (Seahorse Bioscience, MA, USA). Measurements were performed under basal conditions and after the addition of 2 μM oligomycin, 0.5 μM (A549)/0.2 μM (H23) carbonyl cyanide-4-(trifluoromethoxy)phenylhydrazone (FCCP) and 10 μM antimycin. If indicated, 40 μM etomoxir was added (Sigma). OCR was normalized to protein content.

### Mitotracker analysis with flow cytometry

Cells were treated for 24 hours, washed with PBS and then incubated with 100 nM Mitotracker Green (Thermo Scientific, Waltham, MA, USA) in serum-free media for 30 minutes. Thereafter, cells were washed with PBS and measured with flow cytometry (CytoFlex, BeckmanCoulter, Krefeld, Germany) in starvation media. Cells were gated for live and single cells.

### Confocal microscopy of mitochondria

2×10^4^ cells were plated on 8-well chamber slides with glass bottom (Ibidi, Gräfelfing, Germany) and treated for 24 hours with non-starvation or starvation media, washed with PBS and incubated with 100 nM Mitotracker Green (Thermo Scientific) in RPMI SILAC supplemented with arginine and lysine for 30 minutes. Therafter, cells were washed with PBS and reincubated in the respective treatment medium. Immediately, images of representative cells were taken by Nikon A1 confocal microscope (Vienna, Austria). Images were analyzed by ImageJ (National Institutes of Health (NIH)) with the MiNa plugin (Valente *et al*, 2017).

### Detection of ROS

For detection of intracellular mitochondrial superoxide, cells were plated and treated after 24 hours with non-starvation or starvation media. Then cells were washed with PBS, trypsinized and incubated with 2.5 μM of MitoSox (Thermo Scientific) for 20 minutes at 37°C. After washing with PBS, cells were resuspended in PBS and analyzed by flow cytometry (CytoFlex, BeckmanCoulter, Krefeld, Germany). To determine general ROS, cells were treated as described above and ROS were determined with 10 μM of CM-H2DCFDA (Thermo Scientific) upon incubation for 30 minutes at 37°C followed by flow cytometry. Lipid peroxidation was measured in cells treated with or without RSL3 (Selleck chemicals, Houston, TX, USA) with 2 μM Bodipy-C11 dye diluted in HBSS, followed by flow cytometry. Cells were gated for live and single cells.

### Detection of GSH/GSSG

For detection of the GSH/GSSG ratio, cells were plated and treated for 24 hours with non-starvation or starvation media. Then cells were washed with cold PBS and lysed with mammalian cell lysis buffer (Abcam, Cambridge, UK). GSH/GSSG ratio was measured with a commercially available kit (Abcam), according to the manufacturer’s instructions. Samples were deproteinized before measurement.

### Colony formation assay

Cells were plated at the indicated densities onto 6-well plates. 24 hours after plating, cells were washed twice with PBS and incubated in starvation or nonstarvation media for 72 hours, which was followed by a recovery period in normal growth medium. Antioxidants or pro-oxidants were added during the treatment period at the indicated concentrations. After the recovery period, cells were washed with PBS, fixed in methanol:acetic acid (3:1 v/v) and stained using 0.4 % crystal violet. The colony area was determined by using the Colony Area plugin (Guzman *et al*, 2014) and ImageJ software (NIH).

### Cell counting

1×10^5^ (H23) or 7.5×10^4^ (A549) cells were plated and treated for 24 hours with non-starvation or starvation medium and different concentrations of stabilized H_2_O_2_ (Roth, Karlsruhe, Germany). BSO (Sigma) was administered for 72 hours according the duration of treatment in the colony formation experiments. Then cells were trypsinized and counted with the Casy-TT cell counter (Roche Innovatis, Bielefeld, Germany).

### Proliferation

Cells were plated and treated 24 hours with non-starvation or starvation supplemented with 10 mM lactate. For detection of proliferative cells, the EdU Click-iT kit (Thermo Scientific) was utilized according to the manufacturer. In brief, EdU was added at a concentration of 10 μM and incubated for 1.5 hours. Cells were collected, fixed, permeabilized and incubated with the freshly prepared Click iT reaction cocktail. EdU-positive cells were determined by FACS after gating for live and single cells.

### Western blot

Protein was extracted with RIPA buffer (Thermo Scientific) and the concentration was determined by BCA assay (Merck, Vienna, Austria). Proteins were separated by sodium dodecyl sulfate-polyacrylamide gel electrophoresis using the Mini-PROTEAN® electrophoresis unit (BioRad, Hercules, CA) and transferred to a PVDF membrane (BioRad). The following antibodies were used: PCK2 (Abcam, ab187145; 1:2000), total OXPHOS antibody cocktail (Abcam, ab 110411, 1:1000), TOM20 (Cell Signaling Technology, #42406, 1:1000) and β-actin (Santa Cruz, sc-47778 and Sigma, A5441). β-actin was used as a loading control.

### Quantitative real-time PCR (RT-qPCR)

A detailed description of RNA extraction, cDNA synthesis and RT-qPCR and a list of all utilized primers can be found in Appendix Methods.

### Statistics

Data were analyzed with Microsoft Excel 2016 or SPSS, version 23.0 (Chicago, IL, USA). Group comparisons were made using two-sided, unpaired Student’s t-tests, one-group Student’s t-tests or one-way ANOVA with Dunnett post-hoc analysis as applicable, using data from at least three independent experiments. A p<0.05 was considered significant.

## Acknowledgements

We thank A. Bertsch and T. Haitzmann (Medical University of Graz) and D. Broekaert (VIB-KU Leuven) for excellent technical support and we are grateful for the valuable advice by G. Höfler and T. Madl (Medical University of Graz). The study was supported by the Austrian Science Fund (P 28692-B31 to K.L.). G.G. received funding from a DOC Fellowship of the Austrian Academy of Sciences (25282). The study was further supported by DK-MCD W1226 (to W.F.G.), by the Austrian Science Fund Erwin Schroedinger Abroad Fellowship (J4205-B27 to C.T.M.), and by MEFO Graz (to K.L. and W.F.G.). S.-M.F. acknowledges funding from the European Research Council under the ERC Consolidator Grant Agreement n. 771486-MetaRegulation, FWO – Research Projects, KU Leuven – Methusalem Co-Funding and Fonds Baillet Latour.

## Author contributions

G.G, C.T.M., W.F.G., S.-M.F., H.O. and K.L. designed research; G.G., M.P., C.T.M., A.H. and K.L. performed research; G.G., M.P., C.T.M., W.F.G., S.M.F., and. K.L. analyzed data; G.G. and K.L. wrote the paper.

## Conflict of interest statement

The authors declare no conflict of interest.

